# Systems biology of cold adaptation in the polyextremophilic red alga *Galdieria sulphuraria*

**DOI:** 10.1101/565861

**Authors:** Alessandro W. Rossoni, Andreas P.M. Weber

## Abstract

Rapid fluctuation of environmental conditions can impose severe stress upon living organisms. Surviving such episodes of stress requires a rapid acclimation response, e.g., by transcriptional and post-transcriptional mechanisms. Persistent change of the environmental context, however, requires longer-term adaptation at the genetic level. Fast-growing unicellular aquatic eukaryotes enable analysis of adaptive responses at the genetic level in a laboratory setting. In this study, we applied continuous cold stress (28°C) to the thermoacidophile red alga *G. sulphuraria,* which is 14°C below its optimal growth temperature of 42°C. Cold stress was applied for more than 100 generations to identify components that are critical for conferring thermal adaptation. After cold exposure for more than 100 generations, the cold-adapted samples grew ~30% faster than the starting population. Whole-genome sequencing revealed 757 variants located on 429 genes (6.1% of the transcriptome) encoding molecular functions involved in cell cycle regulation, gene regulation, signaling, morphogenesis, microtubule nucleation, and transmembrane transport. CpG islands located in the intergenic region accumulated a significant number of variants, which is likely a sign of epigenetic remodeling. We present 20 candidate genes and three putative cis-regulatory elements with various functions most affected by temperature. Our work shows that natural selection towards temperature tolerance is a complex systems biology problem that involves gradual reprogramming of an intricate gene network and deeply nested regulators.

## Introduction

Small changes in average global temperature significantly affect the species composition of ecosystems. Indeed, 252 Ma years ago up to ~95% of marine species and ~70% of terrestrial vertebrates ceased to exist (Benton, 2008; Sahney and Benton, 2008). This event, known as the Permian-Triassic extinction, was triggered by a sharp increase in worldwide temperature (+8°C) and CO2 concentrations (+2000 ppm) during a period spanning 48,000–60,000 years (McElwain and Punyasena, 2007; Shen et al., 2011; Burgess et al., 2014). In comparison, atmospheric CO2 has increased by ~100 ppm and the global mean surface temperature by ~1°C since the sinking of the Titanic in 1912, a little more than 100 years ago. Anthropogenic climate change and its consequences have become a major evolutionary selective force (Palumbi, 2001). Higher temperatures and CO2 concentrations result in increased seawater acidity, increased UV radiation, and changes in oceanwide water circulation and upwelling patterns. These rapid changes represent dramatically accelerating shifts in the demography and number of species, leading to loss of habitats and biodiversity (Hendry and Kinnison, 1999; Stockwell et al., 2003). A global wave of mass extinction appears inevitable (Kolbert, 2014). In this context, it is relevant to assess the effects of temperature change on genome evolution. Aquatic unicellular eukaryotes are particularly well-suited to addressing this question due to their short generation time and straightforward temperature control of their growth environment.

Microorganisms rapidly acclimate and subsequently adapt to environmental change (López-Rodas et al., 2009; Huertas et al., 2011; Romero-Lopez et al., 2012; Osundeko et al., 2014; Foflonker et al., 2018). These adaptations are driven by natural selection and involve quantitative changes in allele frequencies and phenotype within a short period of time, a phenomenon known as microevolution. The *Galdieria* lineage comprise a monophyletic clade of polyextremophilic, unicellular red algae (Rhodophyta) that thrive in acidic and thermal habitats worldwide (e.g., volcanoes, geysers, acid mining sites, acid rivers, urban wastewaters, and geothermal plants) where they represent up to 90% of the total biomass, competing with specialized Bacteria and Archaea (Seckbach, 1972; Castenholz and McDermott, 2010). Accordingly, members of the *Galdieria* lineage can cope with extremely low pH values, temperatures above 50°C, and high salt and toxic heavy metal ion concentrations (Doemel and Brock, 1971; Castenholz and McDermott, 2010; Reeb and Bhattacharya, 2010; Hsieh et al., 2018). Some members of this lineage also occur in more temperate environments (Gross et al., 2002; Ciniglia et al., 2004; Qiu et al., 2013; Barcytė et al., 2018; Iovinella et al., 2018).

Our work systematically analyzed the impact of prolonged exposure to suboptimal (28°C) and optimal (42°C) growth temperatures on the systems biology of *Galdieria sulphuraria* for a period spanning more than 100 generations. We chose *Galdieria sulphuraria* as the model organism for this experiment due to its highly streamlined haploid genome (14 Mb, 6800 genes) that evolved out of two phases of strong selection for genome miniaturization (Qiu et al., 2015). In this genomic context, we expected maximal physiological effects of novel mutations, thus possibly reducing the fraction of random neutral mutations. Furthermore, we expected a smaller degree of phenotypical plasticity and hence a more rapid manifestation of adaptation at the genetic level.

## Materials and methods

### Experimental design and sampling

A starting culture of *Galdieria sulphuraria* strain RT22 adapted to growth at 42°C was split into two batches, which were grown separately at 42°C (control condition) and 28°C (temperature stress) for a period spanning 8 months. Bacteria were cultured on agar plates under non-photosynthetic conditions, with glucose (50 mM) as the sole carbon source. To select for fast-growing populations, the five largest colonies of each generation were picked. The samples were propagated across generations by iteratively picking the five biggest colonies from each plate and transferring them to a new plate. The picked colonies were diluted in 1 ml Allen Medium containing 25 mmol glucose. The OD750 of the cell suspensions was measured at each re-plating step using a spectrophotometer. Approximately 1,000 cells were streaked on new plates to start the new generation. The remaining cell material was stored at −80°C until DNA extraction. This process was reiterated whenever new colonies with a diameter of 3 mm–5 mm became visible. During the 240 days of this experiment, a total of 181 generations of *Galdieria sulphuraria* RT22 were obtained for the culture grown at 42°C, whereas 102 generations were obtained for *Galdieria sulphuraria* RT22 grown at 28°C.

### DNA extraction and sequencing

DNA from each sample was extracted using the Genomic-tip 20/G column (QIAGEN, Hilden, Germany), following the steps of the yeast DNA extraction protocol provided by the manufacturer. DNA size and quality were assessed via gel electrophoresis and Nanodrop spectrophotometry (Thermo Fisher, Waltham, MA, USA). TruSeq DNA PCR-Free libraries (insert size = 350 bp) were generated. The samples were quantified using the KAPA library quantification kit, quality controlled using a 2100 Bioanalyzer (Agilent, Santa Clara, CA, USA), and sequenced on an Illumina (San Diego, CA, USA) HiSeq 3000 in paired-end mode (1×150bp) at the Genomics and Transcriptomics Laboratory of the Biologisch-Medizinisches Forschungszentrum in Düsseldorf, Germany. The raw sequence reads are retrievable from the NCBI’s Small Read Archive (SRA) database (Project ID: PRJNA513153).

### Read mapping and variant calling

Single nucleotide polymorphisms (SNPs) and insertions/deletions (InDels) were called separately on the dataset using the GATK software version 3.6-0-g89b7209 (McKenna et al., 2010). The analysis was performed according to GATKs best practices protocols (DePristo et al., 2011; Van der Auwera et al., 2013). The untrimmed raw DNA-Seq reads of each sample were mapped onto the genome of *Galdieria sulphuraria* RT22 (NCBI, SAMN10666930) using the BWA aligner (Li and Durbin, 2009) with the −M option activated to mark shorter split hits as secondary. Duplicates were marked using Picard tools (http://broadinstitute.github.io/picard). A set of known variants was bootstrapped for *Galdieria sulphuraria* RT22 to build the covariation model and estimate empirical base qualities (base quality score recalibration). The bootstrapping process was iterated three times until convergence was reached (no substantial changes in the effect of recalibration between iterations were observed, indicating that the produced set of known sites adequately masked the true variation in the data). Finally, the recalibration model was built upon the final samples to capture the maximum number of variable sites. Variants were called using the haplotype caller in discovery mode with -ploidy set to 1 *(Galdieria* is haploid) and −mbq set to 20 (minimal required Phred score) and annotated using snpEff v4.3i (Cingolani et al., 2012). The called variants were filtered separately for SNPs and InDels using the parameters recommended by GATK (SNPs: “QD < 1.0 || FS > 30.0 || MQ < 45.0 || SOR > 9 || MQRankSum < −4.0 || ReadPosRankSum < −10.0”, InDels: “QD < 1.0 || FS > 200.0 || MQ < 45.0 || MQRankSum < – 6.5 || ReadPosRankSum < −10.0”).

### Evolutionary pattern analysis

A main goal of this analysis was implementation of a method that enabled discrimination between random variants and variants that may be connected to temperature stress (non-random variants). The following logic was implemented: All variants were transformed to binary code with regards to their haplotype towards the reference genome. When the haplotype was identical to the reference genome, “0” was assigned. Variant haplotypes were assigned “1”. Random variants were gained and lost without respect to the sampling succession along the timeline and the different temperature conditions. Consequently, a “fuzzy” pattern of, e.g., “110011|0000”, would indicate a mutation between T0 and T1 in the samples taken at 28°C, which was lost in T3 and regained after T5. The binary sequence represents the ten samples, six “cold” and four “warm”, according to their condition (“cold | warm”) and time point of sampling (“28°C_1, 28°C _2, 28°C _3, 28°C _4, 28°C _5, 28°C _6 | 42°C_1, 42°C _3, 42°C _6, 42°C _9”). Hence, the first six digits denote samples taken at 28°C, the latter four digits those taken at 42°C; “000000|0101” would represent a mutation in the T2 sample taken at 42°C that was lost in T3 and regained in T4, and “011010|0101” would represent a variant that does neither with respect to the sampling succession (repeated gain and loss) nor the growth condition of the samples (mutation occurs at both temperatures). By contrast, variants that were gained and fixed in the subsequent samples of a certain growth condition were considered as “non-random variants” that may reflect significant evolutionary patterns. Thus, “111111|0000” would indicate that a mutation between T0 and T1 in the samples taken at 28°C was fixed over the measured period. Similarly, “000000|0111” would indicate a mutation between T2 and T3 in the samples taken at 42°C that was fixed throughout the generations. As such, it was possible to determine all possible pattern combinations for non-random evolutionary patterns. The binary sequence “111111|1111” represented the case where all ten samples contained a different haplotype when compared to the reference genome. In this specific case, systematic discrepancies between the reference genome and the DNA-Seq reads are the cause of this pattern. Variants following the “111111|1111” pattern were removed from the dataset.

### Data accession

The DNA sequencing results are described in Supplementary Table S1. The Illumina HiSeq3000 raw reads reported in this project have been submitted to the NCBI’s Sequence Read Archive (SRA) and are retrievable (FASTQ file format) via BioProject PRJNA513153 and BioSamples SAMN10697271 - SAMN10697280.

### Statistical analysis

Various statistical methods were applied for the different analyses performed in this project. Culture growth was measured for at 28°C (n = 6) and at 42°C (n = 10). Both datasets failed the Shapiro-Wilk normality test (p > 0.05) and showed a visible trend over time. The difference in growth between the poulations was tested using the Wilcoxon rank sum test. Further, timepoints along a timeline constitute a dependent sampling approach by which the growth performance of an earlier timepoint is likely to influence the the growth perfromance of a later timepoint.

Trends in growth over the period of this experiment were tested for significance using Jonckheere-Terpstra’s test for trends. Enrichment of GO categories as well as *k*-mer enrichment was tested using Fisher’s exact test for categorical data, corrected for multiple testing according to Benjamini-Hochberg. The contingency table was set up in such way that the number of times a specific GO was affected by variants was compared with the number of times the same GO was not affected by variants. This category was compared againts the “background” consisting of all other GOs affected by variants and all other unaffected GOs. The same methodology was applied for *k*-mer enrichment testing. Differential gene expression based on previously collected data (Rossoni et al., 2018) was calculated with EdgeR (Robinson et al., 2010) implementing the QLF-test in order to address the dispersion uncertainty for each gene (Lun et al., 2016). All samples taken at 28°C were compared against all samples taken at 42°C/46°C.

## Results

### Culture growth

Samples grown at 42°C were re-plated 10 times during the 7 months of the experiment due to faster colony growth, whereas cultures growing at 28°C were re-plated only six times (**Figure 1A**). Cultures grown at 42°C achieved an average doubling time of 1.32 days, equivalent to an average growth rate of 0.81/day. Cultures grown at 28°C had an average doubling time of 2.70 days, equivalent to an average growth rate of 0.39/day. This difference in doubling time/growth rate between 28°C and 42°C was significant (non-normal distribution of growth rates, Wilcoxon rank sum test, p = 0.0002) (**Figure 1B**). The growth rates reported here were slightly lower than in liquid batch cultures, where growth rates of 0.9/day–1.1/day were measured for heterotrophic cultures grown at 42°C (unpublished data). The changes in growth rate over time were also compared using linear regression (**Figure 1C**). Although the linear regression appears to indicate increasing doubling times in samples grown at 42°C, Jonckheere’s test for trends revealed no significant trend in this dataset (Jonckheere-Terpstra, *p* > 0.05). By contrast, samples grown at 28°C gradually adapted to the colder environment and significantly (Jonckheere-Terpstra, *p* < 0.05) decreased their doubling time by ~30% during the measured period.

**Figure 1.**
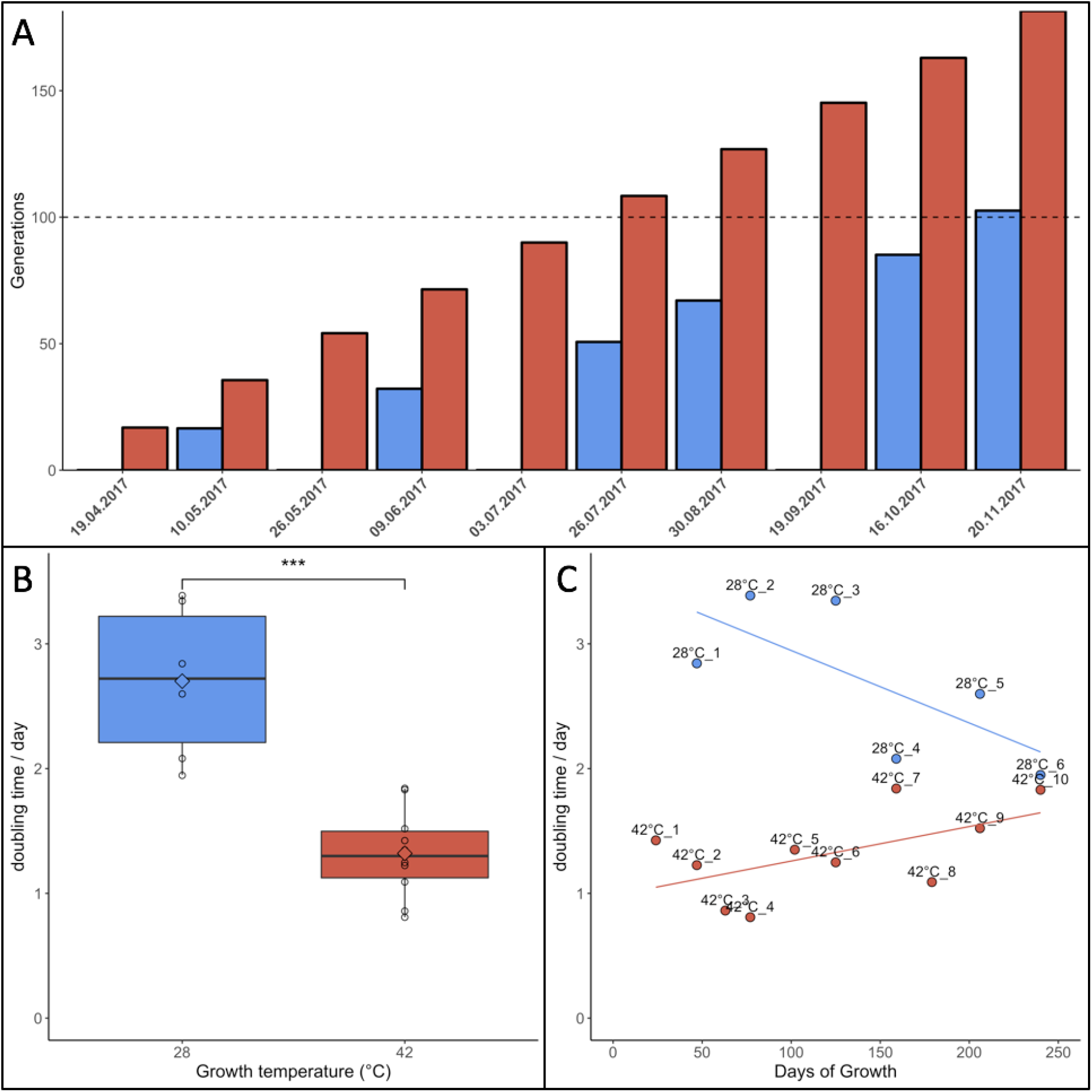
Growth parameters of *Galdieria sulphuraria* RT22. **A**: Cultures were grown heterotrophically at 28°C (blue) and at 42°C (red) on plates made of 1.5% Gelrite mixed 1:1 with 2× Allen Medium containing 50 mM glucose. The fastest growing colonies were iteratively selected and re-plated over a period of ~7 months until >100 generations were achieved under both conditions. Propagation occurred through picking the five biggest colonies from each plate and transferring them to a new plate. **B**: The doubling time at 42°C was 1.32 days on average. The doubling time at 28°C was 2.70 days on average. The differences in growth between 42°C and 28°C were significant (Wilcoxon rank sum test, p = 0.0002). Cultures grown at 42°C were re-plated 10 times due to faster growth. In comparison, cultures growing at 28°C were re-plated only six times. **C**: Samples grown at 42°C grew slightly slower over time. By contrast, samples grown at 28°C appeared to decrease their doubling time. While no statistically significant trend could be detected at 42°C, Jonckheere’s test for trends reported a significant trend towards faster growth for the populations grown at 28°C.

### Variant calling

A total of 470,680,304 paired-end DNA-Seq reads were generated on an Illumina HiSeq 3000 sequencer. Of these, 462,869,014 were aligned to the genome (98.30%) using BWA (**Supplementary Table S1**). The average concordant alignment rate was 99.71%. The average genome coverage was 444× (min = 294×, max = 579×). At least 95.5% of the sequence was covered with a depth of >20×. GATK’s haplotype caller algorithm reported 6,360 raw SNPs and 5,600 raw InDels. The SNPs and InDels were filtered separately according to GATK’s best practice recommendations. A total of 1,864 SNPs and 2,032 InDels passed the filtering process. On average, one SNP occurs every 16,177 nt and one InDel every 44,394 nt. Overall, 66.17% of the filtered variants (2578/3896) were classified as background mutations being at variance with the genome reference (“111111|1111”). The 1243 remaining variants (966 SNPs + 277 InDels) were sorted according to their evolutionary patterns, here called “Random”, “Hot”, and “Cold” (**Figure 2**); 486/1243 (36.5%) are located in the intergenic region and the other 757/1243 (63.5%) in the genic region, including 5’UTR, 3’UTR, and introns. In addition, 1202/1243 (96.7%) variants followed random gain and loss patterns that do not exhibit relevant evolutionary trajectories (**Figure 2**). The remaining 41 variants were gained and fixed over time, thus representing non-random, evolutionary relevant variants. Twenty-three variants were fixed at 28°C and 18 variants at 42°C. Consequently, 23 variants (1.9%) were attributed to the “Cold” pattern (11 intergenic, 12 genic) and 18 variants to the “Hot” pattern (1.4%).

**Figure 2.**
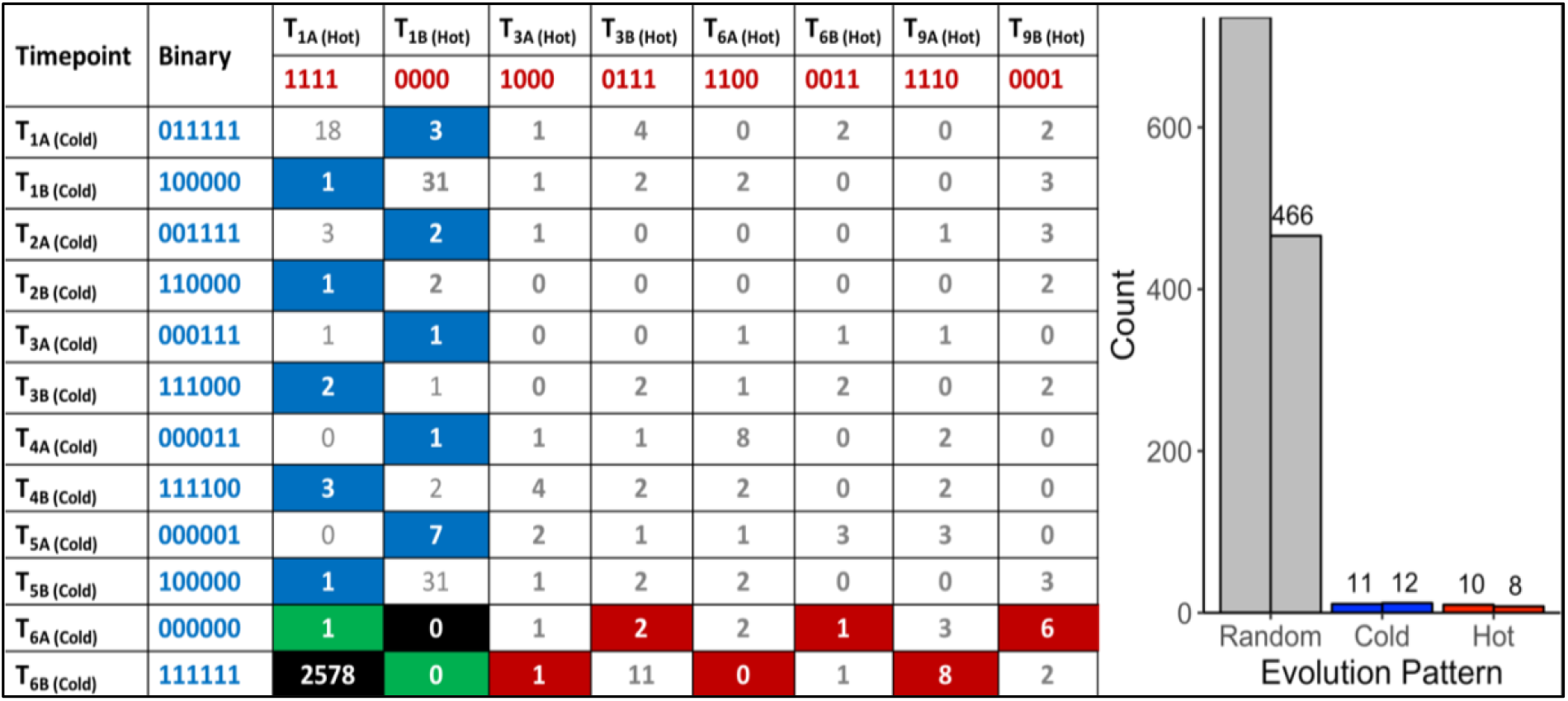
“Random” and “Non-random” variant acquisition patterns. “Non-random” variants were defined as mutations gained at some point during growth at 42°C or 28°C, and fixed in the genome of *G. sulphuraria RT22* during the remaining time points. All variants were translated into binary code according to their haplotype relative to the reference genome. “Cold”: variant was obtained and fixed at 28°C. “Hot”: variant was obtained and fixed at 42°C. **Left**: Evolutionary patterns and their frequencies. In the specific case of patterns “111111|0000” and “000000|1111”, a variant was already gained before the first sampling time point. Hence, it was not possible to determine the condition at which the variant was gained. “Background” mutations represent the cases where the sequence of all samples was in disagreement with the reference genome “111111|1111”. The remaining combinations were considered as “random” evolution patterns. Here, variants were gained but not fixated in the subsequent samples of the same growth condition. The numbers in the boxes indicate the count of a specific pattern. **Right**: Count by variant type. The right column of each category indicates the number of variants located in the intergenic space. The left column counts the number of variants located in the coding sequence.

### GO enrichment-based overview of cellular functions most affected by variants

The vast majority of the 757 genic variants was not fixed over time and did not follow consistent evolutionary patterns (**Figure 2**). However, the frequency at which genes and gene functions were affected by mutation can serve as an indicator of the physiological processes most affected by evolutionary pressure at 28°C and 42°C. Here, we analyzed the functional annotations of the 757 variants located on 429 genes (6.1% of the transcriptome) using GO-Term (GO) enrichment analysis. A total of 1602 unfiltered GOs were found within the genes affected by variants (27.3% of all GOs in *G. sulphuraria* RT22), of which 1116 were found at least twice in the variant dataset. Of those, 234 of the GOs were significantly enriched (categorical data, “native” *vs.* “HGT”, Fisher’s exact test, Benjamini-Hochberg, p ≤ 0.05).

To contextualize the function in broader categories, we manually sorted all significantly enriched GOs into the following ten categories: “Cell Cycle”, “Cytoskeleton”, “Gene Regulation”, “Membrane”, “Metabolism”, “Photosynthesis”, “Stress/Signaling”, “Transport”, “Other”, and “NA” (**Figure 3A**). GO terms belonging to the “NA” category were considered meaningless and excluded from the analysis (e.g., “cell part” [GO:0044464], “biological process” [GO:0008150], “binding” [GO:0005488], “ligase activity” [GO:0016874]).

#### Cell cycle

Functions related to the cell cycle accounted for 79/234 (**33.8%**) of the enriched GOs. Mitosis was affected at every stage: initiation (e.g., “positive regulation of cell proliferation”, GO:0008284, *p* = 0.0024; “re-entry into mitotic cell cycle”, GO:0000320, *p* = 0.0405), DNA replication (e.g., “DNA replication, removal of RNA primer”, GO:0043137, p = 0.0001; “ATP-dependent 5’-3’ DNA helicase activity”, GO:0043141, p = 0.014), prophase (“preprophase band”, GO:0009574, p = 0.0270), metaphase (e.g., “attachment of mitotic spindle microtubules to kinetochore”, GO:0051315, p = 0.0142), anaphase (e.g., “mitotic chromosome movement towards spindle pole”, GO:0007079, p = 0.0134), and telophase (“midbody”, GO:0030496, p = 0.0404). Mutations also accumulated in genes controlling cell cycle checkpoints of mitosis (e.g., “positive regulation of mitotic metaphase/anaphase transition, GO:0045842, p = 0.0270; “mitotic spindle assembly checkpoint”, GO:0007094, p = 0.0441).

Genes with functions involved in cell differentiation and maturation of *Galdieria* were also affected significantly by microevolution during organellogenesis (e.g., “regulation of auxin mediated signaling pathway”, GO:0010928, p = 0.0012; “phragmoplast”, GO:0009524, p = 0.0405; “xylem and phloem pattern formation”, GO:0010051, 0.0012), cell polarity (e.g., “establishment or maintenance of epithelial cell apical/basal polarity”, GO:0045197, p = 0.0012; “growth cone”, GO:0030426, p = 0.0096), and subcellular compartmentalization and localization (“Golgi ribbon formation”, GO:0090161, p = 0.0404; ‘‘establishment of protein localization”, GO:0045184, p = 0.0124). Interestingly, some transcriptional regulators of cell growth seem to be conserved across the eukaryotic kingdom. GOs such as “branching involved in open tracheal system development” (GO:0060446, p = 0.0012) and “eye photoreceptor cell development” (GO:0042462, p = 0.0093) were also found, indicating high amino acid sequence similarity within this category. Further, temperature stress altered genes with functions involved in cell death (e.g., “cell fate determination”, GO:0001709, p = 0.0012; “Wnt signalosome”, GO:1990909, p = 0.0405).

#### Gene regulation

Maintenance of steady and balanced reaction rates across cellular systems is essential for cell survival and poses a major challenge when an organism is confronted with changes in temperature (D’Amico et al., 2002). In this context, the second largest category within the enriched GO terms (49/234, **20.9%**) was related to gene regulation. Besides cell cycle control, thermal adaptation and evolution was orchestrated predominantly through mutations in genes involved in controlling the expression profiles of other genes (“gene expression”, GO:0010467, p = 0.0118). Also, we found a significant proportion of mutations affecting genes linked to the epigenetic control of gene expression, which can occur through methylation of DNA (“hypermethylation of CpG island”, GO:0044027, p = 0.0086), as well as modulation of chromatin density and histone interactions that change the accessibility of whole genomic regions to transcription (“H4 histone acetyltransferase activity”, GO:0010485, p = 0.0040) (Jenuwein and Allis, 2001; Bird, 2002). Further, variants may have altered RNA polymerase efficiency (e.g., “RNA polymerase II transcription factor binding”, GO:0001085, p = 0.0020), mRNA processing (e.g., “regulation of RNA splicing”, GO:0043484, p = 0.0025), post-transcriptional silencing (e.g., “RNA interference”, GO:0016246, p = 0.0093) as well as alteration of ribosome structure components (e.g., “structural constituent of ribosome”, GO:0003735, p = 0.0336) and rRNA methylation components (e.g., “rRNA methylation”, GO:0031167, p = 0.0036). In this regard, GO terms linked to posttranslational protein modification were also enriched (“positive regulation of peptidyl-threonine phosphorylation”, GO:0010800, 0.0086; “N-terminal peptidyl-methionine acetylation”, GO:0017196, p = 0.0007).

#### Cytoskeleton

Microtubuli are long polymers of tubulin that are constituents of the cytoskeleton of every eukaryote. They play a central role in intracellular organization, stability, transport, organelle trafficking, and cell division (Brouhard and Rice, 2018). Because they associate spontaneously, microtubular assembly (e.g., “microtubule nucleation”, GO:0007020, p = 0.0039) and disassembly are mostly driven by tubulin concentrations at the beginning and the end of microtubules once a critical microtubule size is reached (Voter and Erickson, 1984). However, the first steps of microtubule assembly are kinetically unfavorable. Cells solve this issue by using γ-tubulin ring complex as a template (e.g., “tubulin complex”, GO:0045298, p = 0.0031). The reaction equilibrium between tubulin polymerization and monomerization is temperature-dependent and requires accurate regulation (e.g., “tau-protein kinase activity”, GO:0050321, 0.0086). Shifting temperatures from 37°C to 25°C leads to massive microtubular dissociation in homoeothermic species (Himes and Detrich, 1989). Additionally, tubulin adaptations towards lower temperatures have been observed at the level of DNA sequence as well as at the epigenetic level in psychrophilic organisms (Detrich et al., 2000). Microtubule metabolism and its role in cellular physiology accounted for 13/234 (5.6%) of the enriched GOs.

#### Membranes and transport

Another major component that is also influenced by temperature is cell integrity with regards to membrane fluidity (5/234 enriched GOs, **2.1%**) and transport (25/234 enriched GOs, **10.7%**). Cell membranes are selectively permeable and vital for compartmentation and electric potential maintenance. In this context, *Galdieria* is able to maintain near-neutral cytosolic pH against a 10^6^-fold H^+^ gradient across its plasma membrane (Gross, 2000). Membranes maintain a critical range of viscosity to be able to incorporate molecules and transport substrates and nutrients. The fluidity of a membrane is mainly determined by its fatty acid composition. Changes in temperature lead to changes in fatty acid composition, which in turn affect hydrophobic interactions as well as the stability and functionality of membrane proteins and proteins anchored to membranes. Here, we measured a significant enrichment in genes with functions connected to membrane lipid bilayers (e.g., “membrane”, GO:0016020, p = 0.0002; “mitochondrial inner membrane”, GO:0005743, p = 0.0023) as well as membrane-associated proteins (e.g., “integral component of membrane”, GO:0016021, p < 0.0001), transporters (e.g., “amino acid transmembrane transporter activity”, GO:0015171, p = 5.79568E-06), and transport functions (e.g., “transmembrane transport”, GO:0055085, p = 0.0001; “cation transport”, GO:0006812, p = 0.0028). Furthermore, temperature imposes significant restrictions to vesicles, which play a central role in molecule trafficking between organelles and in endocytosis. Vesicle formation in particular appears to be affected by temperature (e.g., “vesicle organization”, GO:0016050, p = 0.0012; “clathrin-coated endocytic vesicle membrane”, GO:0030669, p = 0.0025).

#### Stress and signaling

Cell signaling comprises the transformation of information, such as environmental stress, to chemical signals that are propagated and amplified through the system where they contribute to the regulation of various processes (e.g., “response to stress”, GO:0006950, p = 0.0051; “hyperosmotic response”, GO:0006972, p = 0.0039; “ER overload response”, GO:0006983, p = 0. 0040). Here, we found a total of 18/234 GOs (7.7%) derived from genes involved in cell signaling upon which temperature changes appeared to exhibit significant evolutionary pressure driving the accumulation of variants. A broad array of receptors (G-protein coupled, tyrosine kinases, and guanylate cyclases) performs signal transduction through phosphorylation of other proteins and molecules. The signal acceptors, in turn, influence second messengers and further signaling components that affect gene regulation and protein interactions. GO annotations indicate involvement of temperature in genes coding for receptors (e.g., “activation of protein kinase activity”, GO:0032147, p = 4.95227E-05; “protein serine/threonine/tyrosine kinase activity”, GO:0004712, p = 0.00045547; “protein autophosphorylation”, GO:0046777, p = 1.53236E-06) as well as in genes coding for the signal acceptors (“stress-activated protein kinase signaling cascade”, GO:0031098, p = 6.45014E-06; “cellular response to interleukin-3”, GO:0036016, p = 5.05371E-06; “regulation of abscisic acid-activated signaling pathway”, GO:0009787, p = 0.006712687).

#### Metabolism

Maintaining metabolic homeostasis is paramount for organism survival. The efficiency, speed, and equilibrium of metabolic pathways are modulated by enzymes and the specific kinetics of each reaction. Whereas microorganisms are not capable of controlling the amount of free enthalpy in their systems (chemical equilibriums are temperature-dependent, Δ*G* = −*RTlnK*), they are able to actively adjust their metabolic rates by regulating the amount of available enzyme (“Gene Expression”). Passively, mutations can alter enzyme structure, thereby adjusting the affinity of enzymes towards ligands. Variants affecting the genetic code of genes attributed to this category influence a broad variety of metabolic pathways (e.g., “cellular aromatic compound metabolic process”, GO:0006725, p = 0.0030; “amine metabolic process”, GO:0009308, p < 0.0001) in both anabolism (e.g., “peptidoglycan biosynthetic process”, GO:0009252, p = 0.0015; “glycerol biosynthetic process”, GO:0006114, p = 0.0086), and catabolism (e.g., “glycosaminoglycan catabolic process”, GO:0006027, p = 0.0011). In spite of pronounced changes in gene expression of metabolic enzymes during short-term cold stress in *Galdieria sulphuraria* 074W (Rossoni et al., 2018) and *Cyanidioschyzon merolae* 10D (Nikolova et al., 2017), microevolution of genes directly involved in metabolic steps appeared to play a minor role in long-term temperature adaptation (34/234 GOs, **14.5%**).

#### Photosynthesis

The majority of photosynthetic light reactions are catalyzed by enzymes located in the photosynthetic thylakoid membranes. Hence, photosynthesis is based upon temperature-dependent proteins located in temperature-dependent membranes (Yamori et al., 2014). Abnormal temperatures affect the electron transport chain between the various components of the photosynthetic process (Hew et al., 1969). If the electron transport chain between photosystem I (PSI) and photosystem II (PSII) is uncoupled, electrons are transferred from PSI to oxygen instead of PSII. This process is also known as PSII excitation pressure and leads to a boost of reactive oxygen species. Long-term microevolution did not appear to significantly affect the photosynthetic apparatus of *Galdieria sulphuraria* RT22 (3/234, 1.3%), likely because the experiment was performed under heterotrophic conditions in continuous darkness.

### Variant hotspots and non-random genic variants

To further investigate the temperature adaptation of *Galdieria sulphuraria* RT22, we selected candidate genes for closer analysis using two different approaches. First, we assumed that high mutation rates in a specific gene reflect increased selective force upon its function and regulation. To identify potential targets of temperature-dependent microevolution, we searched for “variant hotspots”, here defined as the 99^th^ percentile of genes most affected by variants. We computed variant number-dependent Z-scores for each gene and extracted genes with a Z-Score > 2.575. This procedure led to identification of seven genes, so-called “variant hotspots”, containing at least seven independent variants per gene. Next, we extracted 41 variants that followed non-random evolutionary patterns, here defined as the gain of a variant and its fixation in the subsequent samples that was exclusive to either 28°C or 42°C (**Figure 2**). Eighteen variants followed an evolutionary pattern defined as “Hot” (1.36%) and 23 variants followed an evolutionary pattern defined as “Cold” (1.59%). These non-random evolutionary patterns describe the gain of a variant and its fixation over time either in the 42 °C dataset (“Hot”, e.g., 000000|0001, 000000|0011), or in the 28°C dataset (“Cold”, e.g., 000001|0000, 000011|0000), respectively. The underlying assumption was that this subset represented beneficial mutations. Synonymous variants were removed from further analysis. As a result, we obtained 13 genes that followed non-random evolutionary patterns (16 non-synonymous variants). An individual functional characterization of each gene is contained in the supplementary material (**Supplementary Listing S1A** for “variant hotspots” and **Supplementary Listing S1B** for “non-random” genic variants).

The gene function of the selected temperature-dependent gene candidates broadly replicated the results of the GO enrichment analysis. Here, we found multiple enzymes involved in cell cycle control and signaling, e.g., an oxidase of biogenic tyramine (Gsulp_RT22_67_G1995), an armadillo/beta-catenin repeat family protein (Gsulp_RT22_107_G5273), the GTPase-activating ADP-ribosylation factor ArfGAP2/3 (Gsulp_RT22_82_G3036), and a peptidylprolyl cis-trans isomerase (Gsulp_RT22_64_G1844). Other candidate genes were involved in transcription and translation, e.g., a NAB3/HDMI transcription termination factor (Gsulp_RT22_83_G3136), or in ribosomal biogenesis (Gsulp_RT22_112_G5896, 50S ribosomal subunit) and required cochaperones (Gsulp_RT22_99_G4499, Hsp40). Three candidate genes were solute transporters (Gsulp_RT22_67_G2013, Gsulp_RT22_118_G6841, Gsulp_RT22_67_G1991). Most interestingly, two genes connected to temperature stress were also affected by mutations. An error-prone iota DNA-directed DNA polymerase (01_Gsulp_RT22_79_G2795), which promotes adaptive point mutation as part of the coordinated cellular response to environmental stress, was affected at 28°C (Napolitano et al., 2000; McKenzie et al., 2001), as well as the 2-phosphoglycerate kinase, which catalyzes the first metabolic step of the compatible solute cyclic 2,3-diphosphoglycerate, which increases the optimal growth temperature of hyperthermophile methanogens (Santos and da Costa, 2002; Roberts, 2005).

### HGT candidates are not significantly involved in temperature microevolution

Horizontal gene transfer has facilitated the niche adaptation of *Galdieria* and other microorganisms by providing adaptive advantages (Schonknecht et al., 2013; Schönknecht et al., 2014; Foflonker et al., 2018). Five of the total 54 HGT gene candidates in *Galdieria sulphuraria* RT22 gained variants (Rossoni et al., 2019). We tested whether a more significant proportion of HGT candidates gained variants in comparison to native genes. This was not the case: HGT candidates did not significantly differ from native genes (categorical data, Fisher’s exact test, *p* < 0.05). Of the HGT candidates, only Gsulp_RT22_67_G2013, a bacterial/archaeal APC family amino acid permease potentially involved in the saprophytic lifestyle of *Galdieria sulphuraria,* accumulated a significant number of mutations (12 variants).

### Genes involved in differential expression were not targeted by mutation

We tested if the 6.1% of genes that gained variants were also differentially expressed during a temperature-sensitive RNA-Seq experiment in *Galdieria sulphuraria* 074W, where gene expression was measured at 28°C and 42°C (Rossoni et al., 2018). Of the 6982 sequences encoded by *G. sulphuraria* RT22, 4569 were successfully matched to an ortholog in *Galdieria sulphuraria* 074W (65.4%); 342 were orthologs to a variant-containing gene, representing 79.7% of all genes containing variants in *Galdieria sulphuraria* RT22. The dataset is representative (Wilcoxon rank sum test, Benjamini-Hochberg, p < 0.05, no differences in the distribution of variants per gene due to the sampling size). Based on this result, 36.3% of the variant-containing genes were differentially expressed. By contrast, 40.1% of the genes unaffected by variants were differentially expressed (**Figure 3B**). The difference between the two subsets was not significant (categorical data, Fisher’s exact test, *p* < 0.05). Hence, genes affected by variance during this microevolution experiment did not react more, or less, pronouncedly to fluctuating temperature.

**Figure 3:**
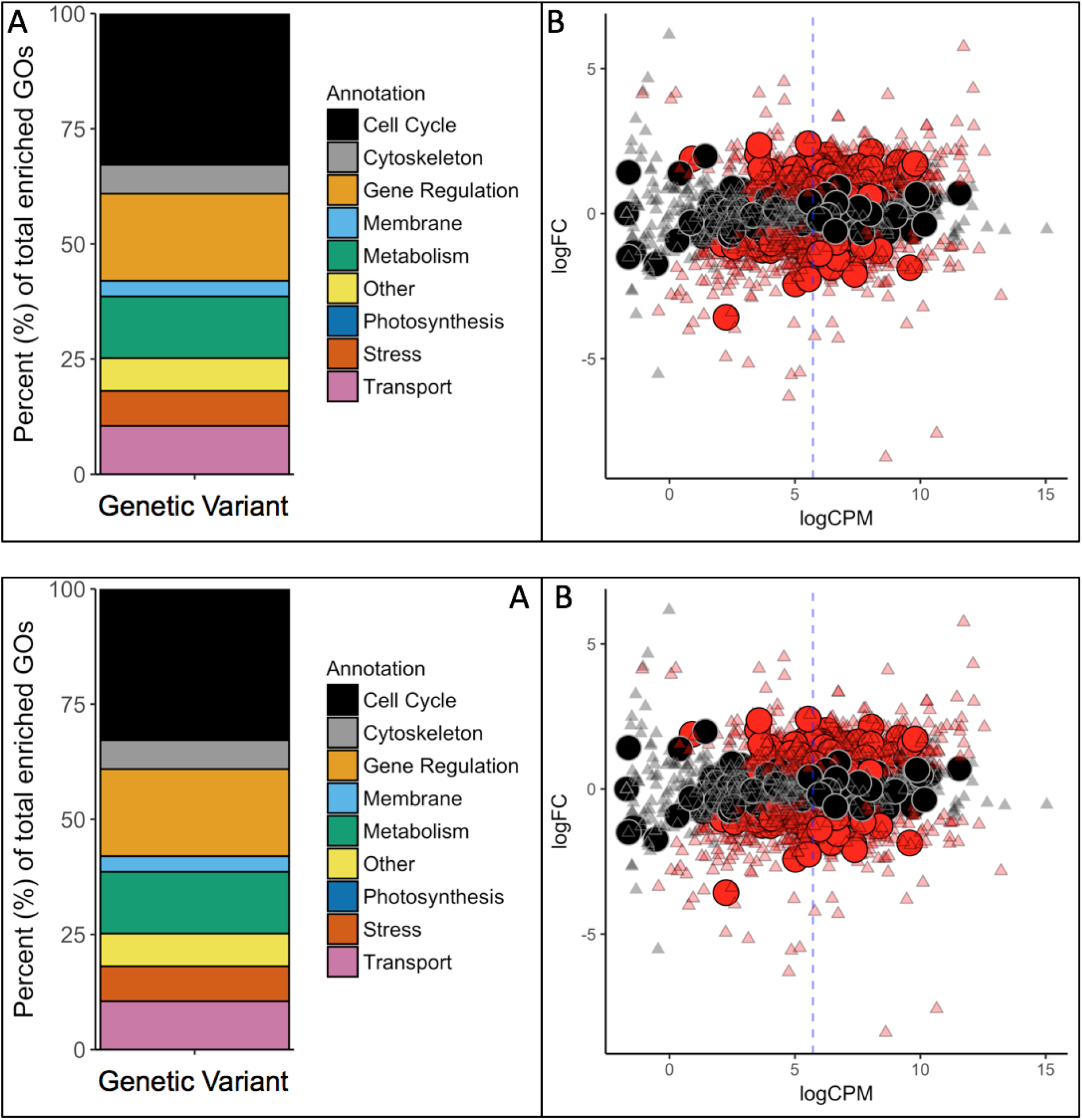
**A:** GO-Term (GO) enrichment analysis revealed the cellular functions most affected by mutations. Each GO was manually revised and attributed to one of the nine categories contained in the legend. **B**: Differential gene expression in *G. sulphuraria* RT22 orthologs in *G. sulphuraria* 074W (reciprocal best blast hit), here measured as log -fold change (logFC) vs. transcription rate (logCPM). Differentially expressed genes are colored red (quasilikelihood F-test, Benjamini-Hochberg, p<= 0.01). Genes affected by variants are shown by large circles. Genes without significant differential expression are represented by triangles. The blue dashes indicate the average logCPM of the dataset. The orthologs in *G. sulphuraria* 074W of genes affected by variance in *G. sulphuraria* RT22 did not show more, or less, differential expression under fluctuating temperature.

### Intergenic variant hotspots

Mutations that affect gene expression strength and pattern are a common target of evolutionary change (Barbosa-Morais et al., 2012). Intergenic DNA encodes cis-regulatory elements, such as promoters and enhancers, that constitute the binding sites of transcription factors and, thus, affect activation and transcriptional rate of genes. Promoters are required for transcriptional initiation but their presence alone results in minimal levels of downstream sequence transcription. Enhancers, which can be located either upstream, downstream, or distant from the genes they regulate, are the main drivers of gene transcription intensity and are often thought to be the critical factors of cis-regulatory divergence (Wray, 2007). Further, epigenetic changes can lead to heritable phenotypic and physiological changes without the alteration of the DNA sequence (Dupont et al., 2009). As a consequence of its evolutionary history, the genome of *Galdieria* sulphuraria is highly deprived of non-functional DNA (Qiu et al., 2015). Here, we performed variant enrichment analysis of the intergenic space based on *k*-mers ranging from *k*-mer length 1 (4 possible combinations, A|C|G|T) to *k*-mer length 10 (1,048,576 possible combinations) to identify intergenic sequence patterns prone to variant accumulation (**Table 1**). The enriched *k*-mers were screened and annotated against the PlantCARE (Lescot et al., 2002) database containing annotations of plant cis-acting regulatory elements. Only partial hits were found, possibly due to the large evolutionary distance between plants and red algae, more specifically the *Galdieria* lineage, which might explain the divergence between *cis*-regulatory sequences (Wittkopp and Kalay, 2012). The sequence “CG”, which is the common denominator of CpG islands (Deaton and Bird, 2011), was found enriched within the *k*-mer set of length 2. In addition, partial hits to the PlantCARE database with a *k*-mer length > 5 were considered as potential hits. Using this threshold, we found three annotated binding motifs, the OCT (octamer-binding motif) (Zhao, 2013), RE1 (Repressor Element 1) (Paonessa et al., 2016), and 3-AF1 (accessory factor binding sites) (Scott et al., 1996; Rhen and Cidlowski, 2005).

**Table 1.**
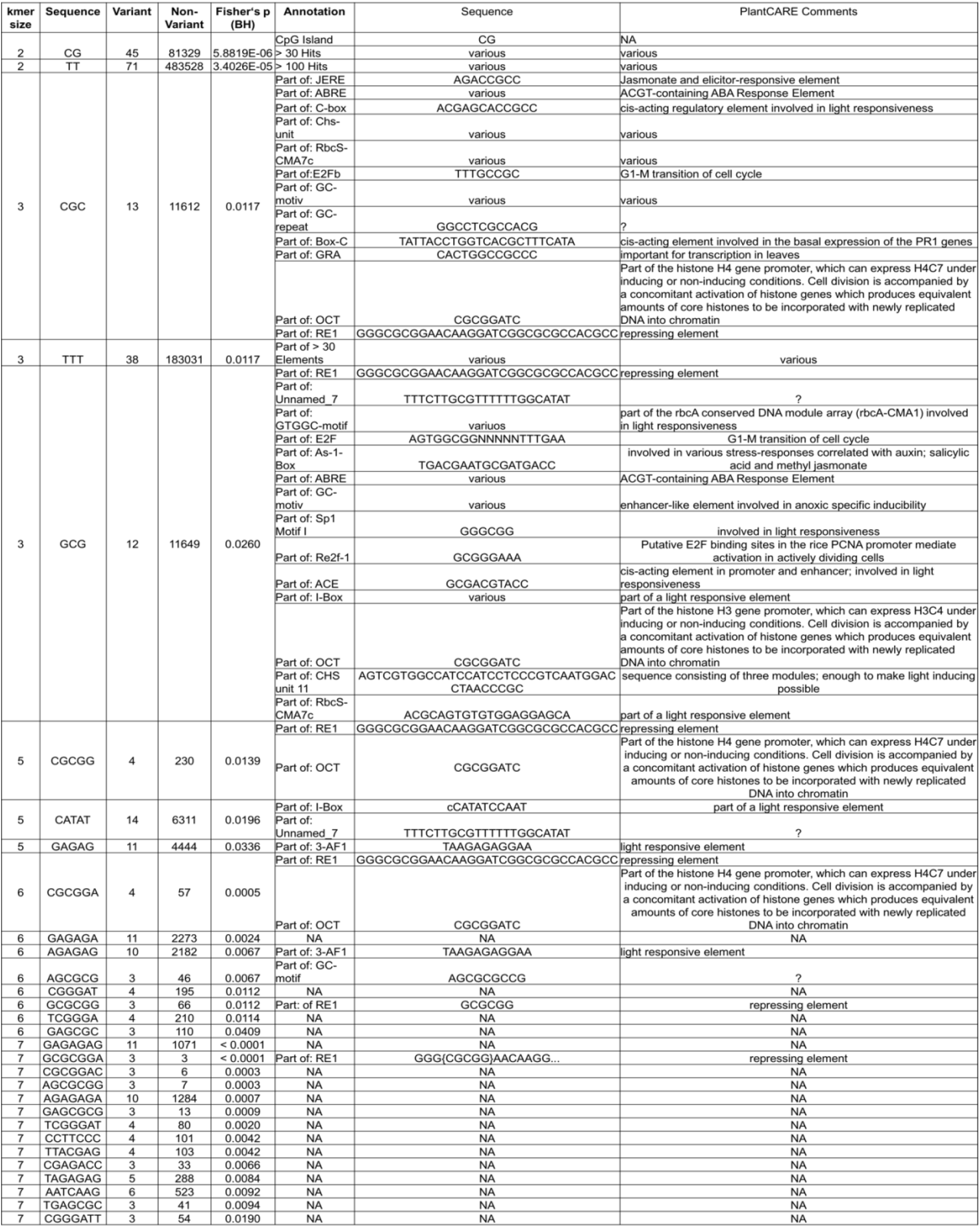
*K*-mer screen of intergenic regions. The non-coding sequence of *Galdieria sulphuraria* RT22 was screened using *k*-mers spanning 1–10 nucleotides. The *k*-mers of each length were tested for variant enrichment (Fisher’s exact test). Only significantly enriched *k*-mers are shown here. *K*-mers longer than eight nucleotides did not produce any database hits and are not shown. **<u>k</u>-mer size**: length of the analyzed *k*-mer. **Sequence**: the sequence of the *k*-mer. **Variant**: Number of *k*-mers with specific sequence affected by variants. **Non-Variant**: Number of *k*-mers with a specific sequence not affected by variants. **Fisher’s p (BH)**: Benjamini-Hochberg post-hoc corrected *p*-value of Fisher’s enrichment test. **Annotation**: PlantCARE identifier (ID) of regulatory element. “Part of:” indicates a partial hit of the k-mer sequence to the database entry. **ID Sequence**: Full sequence of the regulatory element. **PlantCARE Comments**: Additional information provided by PlantCARE.

## Discussion

### Growth rates adapt to temperature

In this study, we subjected two populations of *Galdieria sulphuraria* RT22 to a temperature-dependent microevolution experiment for 7 months. One culture was grown at 28°C, representing cold stress, and a control culture was grown at 42°C. This experiment aimed to uncover the genetic acclimation response to persistent stress, rather than the short-term acclimation response of *Galdieria sulphuraria* to cold stress (Rossoni et al., 2018). We performed genomic re-sequencing along the timeline to measure changes in the genome sequence of *Galdieria sulphuraria* RT22. After 7 months, corresponding to ~170 generations of growth at 42°C and ~100 generations of growth at 28°C, the cold-adapted cultures decreased their doubling time by ~30%. The control cultures maintained constant growth, although a trend to slower growth might occur (**Figure 1**). A similar increase in the growth rate was also observed in the photoautotrophic sister lineage of *Cyanidioschyzon,* where cultures of *Cyanidioschyzon merolae* 10D were grown at 25°C for a period of ~100 days, albeit under photoautotrophic conditions. This study found that the cold-adapted cultures outgrew the control culture at the end of the experiment (Nikolova et al., 2017). While faster doubling times at 28°C can be attributed to gradual adaptation to the suboptimal growth temperatures, we may only speculate about the causes leading to slower growth in the control condition (42°C). Perhaps *Galdieria sulphuraria* RT22, which originated from the Rio Tinto river near Berrocal (Spain), may be able to thrive at high temperatures, but not for such a prolonged period.

### Cultures grown at 28°C accumulate twice the number of mutations as compared to controls

We identified 1243 filtered variants (966 SNPs + 277 InDels), of which 757 (63.5%) were located on the coding sequence of 429 genes and 486 (36.5%) in the intergenic region. The mutation rate was estimated to be 2.17×10^−6^/base/generation for samples grown at 28°C and 1.10×10^−6^/base/generation for samples grown at 42°C, which we interpret as an indication of greater evolutionary stress at 28°C. Hence, suboptimal growth temperatures constitute a significant stress condition and promote the accumulation of mutations. In comparison, mutation rates in other microevolution experiments were 1.53×10^−8^/base/generation-6.67×10’ ^11^/base/generation for the unicellular green freshwater alga *Chlamydomonas reinhardtii* and 5.9×10^−9^/base/generation in the green plant *Arabidopsis thaliana* (Ness et al., 2012; Sung et al., 2012; Perrineau et al., 2014). The 100-fold higher evolutionary rates in comparison to *Chlamydomonas reinhardtii* might result from the selective strategy employed in this experiment (only the five biggest colonies were selected to start the next generations). Although the cold-stressed samples accumulated twice as many mutations per generation in comparison to the control condition, the number of gained variants over the same period was higher in the 42°C cultures due to faster growth rates.

### Cell cycle and transcription factors are the main drivers of temperature adaptation

The impact of temperature-driven microevolution on the cellular functions of *Galdieria sulphuraria* RT22 was analyzed using GO enrichment analysis. More than 75% of the 234 significantly enriched GOs affected genes functions involved in the processes of cell division, cell structure, gene regulation, and signaling. In short, the cellular life cycle appears to be targeted by variation at any stage starting with mitosis, morphogenesis, and finishing with programmed cell death. By contrast, genes directly affecting metabolic processes were less affected by mutation and made up only 10% of the enriched GOs. These observations were also confirmed through the functional annotation of the seven genes most affected by variants (“variant hotspots”) as well as the 13 genes carrying non-synonymous variants with non-random evolutionary patterns.

### The intergenic space in Galdieria is equally affected as coding regions

Historically, intergenic DNA has frequently been considered to represent non-functional DNA. It is now generally accepted that mutations affecting intergenic space can heavily influence the expression intensity and expression patterns of genes. Variants altering the sequence of *cis*-regulatory elements are a common source of evolutionary change (Wittkopp and Kalay, 2012). Due to two phases of genome reduction (Qiu et al., 2015), the genome of *Galdieria* is highly streamlined and the intergenic space accounts for only 36% of its sequence. As a consequence, it is assumed that *G. sulphuraria* lost non-functional intergenic regions that are affected by high random mutation rates in other organisms. In this experiment, variants accumulated proportionally between the genic and the intergenic space, which we interpret as an indication of high relevance of the non-coding regions in *Galdieria.* K-mer analysis revealed significant enrichment of variants occurring in CpG islands. CpG islands heavily influence transcription on the epigenetic level through methylation of the cytosines. In mammals, up to 80% of the cytosines in CpG islands can be methylated, and heavily influence epigenetic gene expression regulation. Furthermore, they represent the most common promotor type in the human genome, affecting transcription of almost all housekeeping genes and the portions of developmental regulator genes (Jabbari and Bernardi, 2004; Saxonov et al., 2006; Zhu et al., 2008). Hence, temperature adaptation is not only modulated through accumulation of mutations in the genetic region but equally driven by the alteration of gene expression through epigenetics and mutations affecting the non-coding region.

## Conclusion

We show here that the significant growth enhancement of samples grown at 28°C over more than 100 generations was driven mainly by mutations in genes involved in the cell cycle, gene regulation, and signal transfer, as well as mutations that occurred in the intergenic regions, possibly changing the epigenetic methylation pattern and altering the binding specificity to *cis*-regulatory elements. Our data indicate the absence of a few specific “key” temperature switches. Rather, it appears that the evolution of temperature tolerance is underpinned by a systems response which requires the gradual adaptation of an intricate gene expression network and deeply nested regulators (transcription factors, signaling cascades, cis-regulatory elements). Our results also emphasize the difference between short-term acclimation and long-term adaptation with regard to temperature stress, highlighting the multiple facets of adaptation that can be measured using different technologies. The short-term stress response of *Galdieria sulphuraria* and the long-term stress response in *Cyanidioschyzon merolae* were quantified using transcriptomic and proteomic approaches, respectively (Nikolova et al., 2017; Rossoni et al., 2018). At the transcriptional and translational levels, both organisms reacted towards maintaining energetic and metabolic homeostasis by increased protein concentrations, adjusting the protein folding machinery, changing degradation pathways, regulating compatible solutes, remodeling of the photosynthetic machinery, and tuning the photosynthetic capacity. SNP and InDel calling revealed underlying regulators mostly affected by variation which are potential drivers of altered transcript and protein concentrations and ultimately determine physiology and phenotype. Some issues, however, remained unresolved. Is the observed growth phenotype permanent, or is it mostly derived from epigenetic modification which could be quickly reversed? We also did not investigate the temperature-dependent differential splicing (Bhattacharya et al., 2018; Qiu et al., 2018) apparatus in *Galdieria,* or the impact of non-coding RNA elements, both of which may provide additional layers for adaptive evolution (van Bakel et al., 2010).

## Acknowledgements

This work was supported by the Deutsche Forschungsgemeinschaft under Germany’s Excellence Strategy – EXC-2048/1 – Project ID: 390686111 and DFG-ANR grant Mo-MiX (WE 2231/21-1). We thank the “Genomics and Transcriptomics laboratory” of the “Biologisch-Medizinischen Forschungszentrum” (BMFZ) at the Heinrich-Heine-University of Düsseldorf (Germany) for technical support and conducting the Illumina sequencing.

