## Supplemental Materials for "Systems biology of cold adaptation in the polyextremophilic red alga *Galdieria sulphuraria*"

### SUPPLEMENTARY MATERIAL

**Supplementary Table S1**: DNA-Seq and mapping statistics. 150x2 bp paired-end reads were generated through Illumina ® HiSeq3000 sequencing and mapped to the genome of *G. sulphuraria* RT22 using bwa. **Sample ID**: Unique ID of the sequenced sample. **Tot. Reads**: Numer of generated reads. **Aligned Reads**: Number of aligned reads. **% Aligned Reads**: Percentage of aligned reads. **Indel rate**: Ratio of InDels per aligned read. **Mismatch rate**: mismatches per read. **Paired Alignment rate**: Read pairs concordantly aligned on the same contig, taking into account their insert size. **Genome Coverage**: average genome per base coverage.

| Sample ID | Tot. Reads | Aligned Reads | % Aligned Reads | Indel rate | Mismatch rate | Paired Alignment rate | Genome Coverage |
|---|---|---|---|---|---|---|---|
| 42°C_9 | 40748234 | 40092215 | 98.39% | 0.000276 | 0.0056 | 0.997197 | 384.85 |
| 42°C_6 | 57701476 | 56813602 | 98.46% | 0.000283 | 0.0058 | 0.996572 | 545.36 |
| 42°C_3 | 47770978 | 47091507 | 98.58% | 0.000229 | 0.0052 | 0.997448 | 452.03 |
| 42°C_1 | 43302404 | 42727881 | 98.67% | 0.00026 | 0.0056 | 0.997356 | 410.15 |
| 28°C_6 | 46621914 | 45959186 | 98.58% | 0.000257 | 0.0054 | 0.997272 | 441.16 |
| 28°C_5 | 47371438 | 46478585 | 98.12% | 0.000267 | 0.0055 | 0.997041 | 446.15 |
| 28°C_4 | 49079250 | 48317389 | 98.45% | 0.000273 | 0.0057 | 0.99693 | 463.80 |
| 28°C_3 | 45309302 | 44473029 | 98.15% | 0.00027 | 0.0057 | 0.996947 | 426.90 |
| 28°C_2 | 31506428 | 30594494 | 97.11% | 0.000264 | 0.0056 | 0.997038 | 293.68 |
| 28°C_1 | 61268880 | 60321126 | 98.45% | 0.000303 | 0.0062 | 0.996813 | 579.02 |

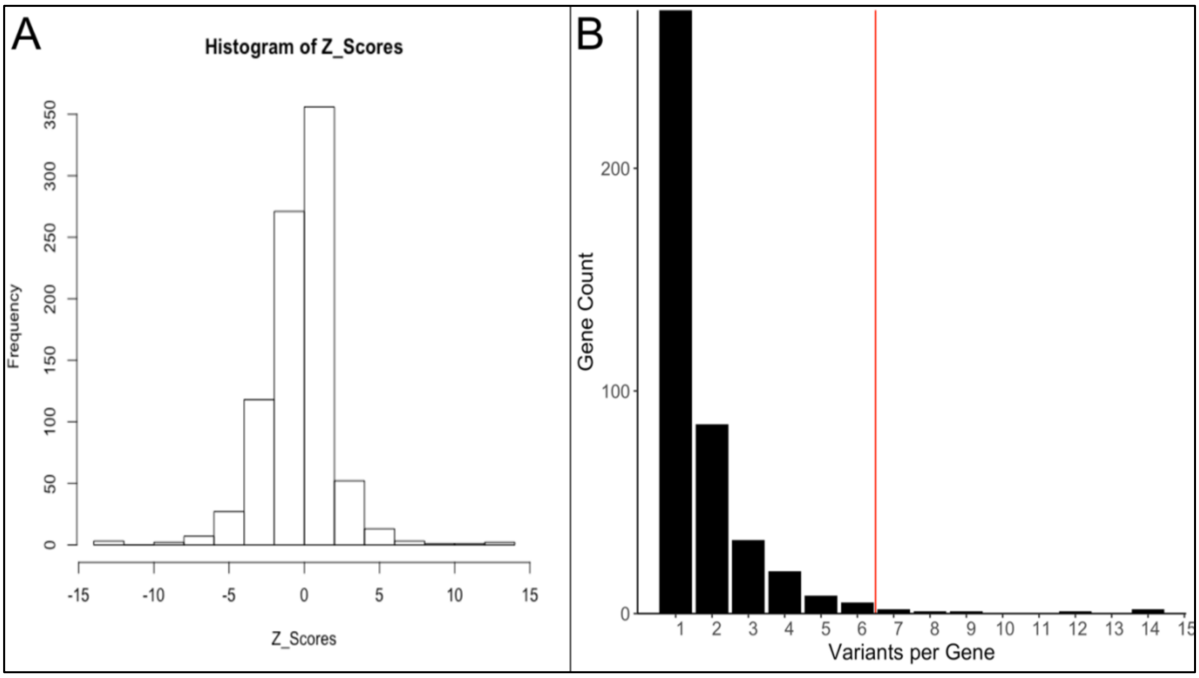


**Supplementary Figure S1** – Z-score distribution of variants per gene. **A:** The calculation of Z-Scores requires a normal distribution of the data in order to calculate standard deviation. To calculate standard deviation, we mirrored the positive distribution of variants per gene by mirroring the number of variants per gene for each gene at x = 0. Doing so, we obtained a normal distribution (Shapiro-Wilk normality test, $p > 0.05$). **B**: Genes on the right side of the red line belong to the 99$^{th}$ percentile of genes with the most variants per gene. Z-scores above 2.56 translated into 7 or more variants per gene.

#### Supplementary Listing S1A - Variant hotspots

*Gsulp_RT22_67_G1995*: (14 variants, Z-score = 5.94) is a tyramine oxidase using copper and 2,4,5-trihydroxyphenylalanine (TPQ) as cofactors which is involved in the metabolism of tyramine, a biogenic amine acting as signal in eukaryotes where it fulfills signaling functions including cell differentiation and growth and detoxification [68] (Signalling, Cell Cycle). Its activation/deactivation is highly regulated through various processes that can target the quinone cofactor or specific amino acid residue of the target enzyme [69].

*Gsulp_RT22_116_G6539*: (14 variants, Z-score = 5.94) is a hypothetical protein which is possibly involved in protein binding (GO:0005515, only available annotation).

*Gsulp_RT22_67_G2013*: (12 variants, Z-score = 5.10) is an APC family amino acid permease involved in the transmembrane transport of amino acids into the cell (Transport) that was potentially acquired via HGT. While temperature affects the maximum uptake rates of amino acids, it does not express a major effect on protein inhibition [70]. Of the 12 variants affecting this gene, only two are synonymous. We observe a trend from bigger to smaller AA (p.Thr394Ser, p.Ile343Val, p.Ile279Val, p.Ser365Gly, p.Ile257Val, p.Thr349Ala) and changes in polarity/charge (p.Asn123Lys, p.Asn123Asp, p.Glu122Gln).

*Gsulp_RT22_107_G5273*: (9 variants, Z-score = 3.82) is an armadillo/beta-catenin repeat family. Armadillo repeats are contained in proteins such as catenin, importin and plakoglobin, and others [71, 72]. β-catenin, the best-characterized protein containing armadillo repeats, is a dual function protein acting as a subunit of the cadherin protein complex (cell-cell adhesion, binding actin filaments across cells) and as an intracellular signal transducer of Wnt signaling (growth control) during embryonic development [73].

*Gsulp_RT22_99_G4499*: (8 variants, Z-score = 3.40) is the molecular chaperone DnaJ, also known as Hsp40. Various Hsp40's act as cochaperones of Hsp70 and regulate its localizations as well as affinity towards complexation with client proteins [74].

*Gsulp_RT22_112_G5896*: (7 variants, Z-score = 2.97) encodes the mitochondrial or chloroplast 50S ribosomal protein L1 involved in the translocation of deacylated tRNAs from the P to the E site [75]. Both the correct biogenesis of ribosomes as well as the process of ribosomal translation are highly temperature dependent and necessitate proper chaperone function, such as DnaJ/Hsp40 [76], which we also found to be a mutation hotspot in this dataset.

*Gsulp_RT22_116_G6590*: (7 variants, Z-score = 2.97) is a LYR family protein. The Nfs1 homolog in yeast encodes for an interacting protein essential in the early steps of Fe/S cluster biogenesis in mitochondria by interacting with Nfs1, the mitochondrial cysteine desulfurase, thus modulating Fe/S cluster assembly [77].

**Supplementary Listing S1B - Further non-random genic variants**

*Gsulp_RT22_82_G3036*: (1 "Cold" variant) has obtained a single intron variant (c.1761+13C>T) which is present in all samples collected at 28°C (28°C_1 – 28°C_6), but absent from all samples taken at 42°C (42°C_1 – 42°C_9). Hence it was obtained within the first 16 generations of growth at cold temperatures. The annotation of this gene suggests the GTPase-activating ADP-ribosylation factor ArfGAP2/3 (Glo3 in yeast, AGD8/9/10 in *A. thaliana*) [78]. ArfGAP2/3 is paramount for regulating the activity amplitude of Arf1 (signaling). Arf1 is a small GTPase central for the generation of vesicles and the correct transport localization of various compounds from the trans-Golgi network to the plasma membrane [79]. In temperature context, Δglo3 mutants in yeast displayed growth defects at 15°C [80]. Unless affecting splice site and other structural components, intronic mutations may affect gene expression levels. Hence, this mutation may add a beneficial transcriptional activity on ArfGAP2/3 which in turn controls the amplitude of Arf1.

*Gsulp_RT22_83_G3136*: (2 "Cold" variant, 4 variants in total) accumulated two intron variants (c.416-19_416-18insG, c.416-22_416-21insT), both measured at the fifth sampling timepoint at 28°C (28°C_5, 67 – 85 generations). Changes in transcription levels of the gene may be assumed. The gene encodes a nuclear polyadenylated RNA-binding NAB3/HDMI protein functioning as transcription termination factor modulating the transcriptional termination at the 3' end of target genes, thus affecting their transcriptional regulation (the underlying mechanism is not fully understood). In yeast, NAB3/HDM1 is also connected to the longevity of CLN3 mRNA [81]. CLN3 is a G1 cyclin that functions as a rate-limiting activator of cell cycle initiation (gene regulation, signaling) which affects the transcription of NAB3/HDM1 thus indirectly influences the temperature dependent cell cycle regulation.

*Gsulp_RT22_64_G1844*: (2 "Cold" variants, 4 variants in total) is a peptidylprolyl cis-trans isomerase (cyclophilin), possibly located in the chloroplast, containing a U box domain [82]. The U box domain indicates that the protein can be posttranslationally modified through targeted ubiquitination. Cyclophilins occur in all subcellular compartments and are involved in processes such as protein maturation by catalyzing protein folding, receptor complex stabilization, receptor signaling, apoptosis, RNA processing and spliceosome assembly (gene regulation, signaling) [83]. Both variants were obtained in the last 28°C timepoint (28°C_6, 86 – 101 generations) and are transversions located in the 5'UTR sequence, one of which (c.-18A>T) leads to the gain of a premature start codon increasing the protein length by 6 amino acids.

*Gsulp_RT22_118_G6841*: (1 "Cold" variant) was annotated as arabinose-proton symporter belonging to the MFS family of sugar transporters, which are not directly affected by temperature (temperature only affects $V_{max}$, not $K_m$) [84]. This gene gained one missense variant (p.Trp209Leu) at the fourth 28°C sampling time point (28°C_4, 50 – 66 generations). While tryptophan is bulkier than leucine, both amino acids are hydrophobic.

*Gsulp_RT22_65_G1905*: (1 "Cold" variant, 3 variants in total) was annotated as inositol-2-dehydrogenase (myo-inositol) catalyzing the oxidation of the carbocyclic myo-inositol [myo-inositol + $NAD^+$ ⇌ 2,4,6/3,5-pentahydroxycyclohexanone + NADH + $H^+$] [85]. It is probably located on clathrin vesicles. Clathrin vesicles are generated by the complete invagination of clathrin-coated pits on the plasma membrane which leads to the formation of cytosolic vesicles. Hence, they play a relevant part in endocytosis, by which extracellular material can be trapped into vesicles and imported into the cell [86]. The temperature relevant mutation is a missense variant (p.Asn277Ser) occurred at the fourth 28°C sampling time point (28°C_4, 50 – 66 generations). Both Asn and Ser are small polar amino acids with uncharged side chains.

*Gsulp_RT22_79_G2795*: (1 "Cold" variant) was annotated as Iota DNA polymerase/DNA polymerase IV which gained a missense variant (p.Pro443Ser) in the last sampling point of 28°C (28°C_6, 86 – 101 generations). It is an error-prone DNA directed DNA polymerase which is activated upon SOS signaling as promoting adaptive point mutations as part of the coordinated cellular response to environmental stress [43, 44]. While proline is a unpolar amino acid with an aliphatic side chain, serine is polar.

*Gsulp_RT22_118_G6798*: (1 "Hot" variant, 3 variants in total) was annotated as P-loop/NTPase domain-containing 2-phosphoglycerate kinase. 2-phosphoglycerate kinase catalyzes the first synthesis step of the unusual compound cyclic 2,3-diphosphoglycerate (cDPG) [87]. cDPG is a compatible solute that has been found increasing the optimal growth temperature of hyperthermophile methanogens as well as salt stress halotolerant microorganism [45, 46]. The variant was measured at the sixth 42°C sampling time point (42°C_6, 54 – 109 generations) and constitutes a missense variant (p.Met747Val).

*Gsulp_RT22_71_G2297*: (1 "Hot" variant) is a hypothetical protein located in membranes that gained an intron variant (c.856+11T>A), possibly influencing its expression pattern, at the third 42°C sampling time point (42°C_3, 16 – 54 generations).

*Gsulp_RT22_84_G3173*: (1 "Hot" variant) is a hypothetical protein located in membranes that gained a missense variant (Arg115Gln) at the sixth 42°C sampling time point (42°C_6, 54 – 109 generations).

*Gsulp_RT22_85_G3288*: **(**1 "Hot" variant**)** gained an in-frame insertion variant (p.Arg302dup) at the ninth 42°C sampling time point (42°C_9, 109 – 162 generations). It was annotated as chloroplastic/mitochondrial Phosphoribosylformylglycinamidine synthase (FGAM synthase) which catalyzes the fourth step of purine biosynthesis [88].

*Gsulp_RT22_67_G1991*: **(**1 "Hot" variant**)** was annotated as a xanthine/uracil/vitamin C permease transmembrane transporter that is potentially subjected do intrinsic misfolding [89]. The majority of members belonging to this family are uncharacterized and potentially transport other substrates. It gained a missense variant (p.Ile584Met) during the ninth 42°C sampling time point (42°C_9, 109 – 162 generations). As temperature does not directly affect transport protein functionality, we suspect an adaptation towards altered membrane viscosity.

*Gsulp_RT22_67_G2027*: (1 "Hot" variant, 4 variants in total) was annotates as the interconvertible, molybdenum cofactor-dependent enzyme xanthine dehydrogenase/oxidase (XDH/XO) which catalyzes the oxidation of hypoxanthine (natural purine derivative) to xanthine and the subsequent oxidation of xanthine to uric acid while generating superoxide and hydrogen peroxides as by-products [90, 91]. The variant (missense, p.Val797Ala) was measured in the ninth 42°C sampling time point (42°C_9, 109 – 162 generations).

*Gsulp_RT22_99_G4476*: (2 "Hot" variants, 3 variants in total) was annotated as γ-tubulin ring complex 3 protein component homologous to Spc98. Spc98 is connected to the recruitment of tubulin at the nuclear surface and possibly along multiple microtubule-nucleating sites in plants [92, 93]. Both variants are frameshift variants that affect the same position (c.2134dupT, c.2133_2134insA). Both lead to a premature stop codon after changing the terminal amino acids to FLERVVK, leading to a loss of trailing 143 amino acids out of its total 2580.

*Gsulp_RT22_107_G5261*: (1 “Cold” variant, 4 variants in total), Gsulp_RT22_79_G2814 (1 “Cold” variant), Gsulp_RT22_104_G4978 (1 “Hot” variant) and Gsulp_RT22_65_G1893 (1 “Hot” variant, 6 variants in total) gained synonymous variants each and were not further considered.
